# *Vibrio cholerae* genomic diversity within and between patients

**DOI:** 10.1101/169292

**Authors:** Inès Levade, Yves Terrat, Jean-Baptiste Leducq, Ana A. Weil, Leslie M. Mayo-Smith, Fahima Chowdhury, Ashraful I. Khan, Jacques Boncy, Josiane Buteau, Louise C. Ivers, Edward T. Ryan, Richelle C. Charles, Stephen B. Calderwood, Firdausi Qadri, Jason B. Harris, Regina C. LaRocque, B. Jesse Shapiro

## Abstract

Cholera is a severe, waterborne diarrheal disease caused by toxin-producing strains of the bacterium *Vibrio cholerae*. Comparative genomics has revealed “waves” of cholera transmission and evolution, in which clones are successively replaced over decades and centuries. However, the extent of *V. cholerae* genetic diversity within an epidemic or even within an individual patient is poorly understood. Here, we characterized *V. cholerae* genomic diversity at a micro-epidemiological level within and between individual patients from Bangladesh and Haiti. To capture within-patient diversity, we isolated multiple (8 to 20) *V. cholerae* colonies from each of eight patients, sequenced their genomes and identified point mutations and gene gain/loss events. We found limited but detectable diversity at the level of point mutations within hosts (zero to three single nucleotide variants within each patient), and comparatively higher gene content variation within hosts (at least one gain/loss event per patient, and up to 103 events in one patient). Much of the gene content variation appeared to be due to gain and loss of phage and plasmids within the *V. cholerae* population, with occasional exchanges between *V. cholerae* and other members of the gut microbiota. We also show that certain intra-host variants have phenotypic consequences. For example, the acquisition of a *Bacteroides* plasmid and nonsynonymous mutations in a sensor histidine kinase gene both reduced biofilm formation, an important trait for environmental survival. Together, our results show that *V. cholerae* is measurably evolving within patients, with possible implications for disease outcomes and transmission dynamics.

**Author Summary:** *Vibrio cholerae* is the etiological agent of cholera, a severe diarrheal disease endemic to Bangladesh and responsible for global outbreaks, including one ongoing in Haiti. Certain bacterial pathogens can evolve and diversify within the human host, often altering virulence and antibiotic resistance. However, most examples of within-host evolution have come from chronic infections, in which the pathogen has sufficient time to mutate and diversify, and little attention has been paid to more acute infections such as the one caused by *V. cholerae*. The goal of this study was to measure the extent of within-host evolution of *V. cholerae* within individual infected patients. By sequencing multiple bacterial isolates_from each of eight patients from Bangladesh and Haiti, we found that cholera patients can harbor a diverse population of *V. cholerae*. As expected for an acute infection, this diversity is limited, ranging from zero to three point mutations (single nucleotide variants) per patient. However, gene gain/loss events are more prevalent than point mutations, occurring in every single patient, and sometimes involving the transfer of dozens of genes on plasmids. Even if rare, point mutations and gene gain/loss events may be maintained by natural selection, and can alter clinically-and environmentally-relevant phenotypes such as biofilm formation. Therefore, within-patient evolution has the potential to impact clinical and epidemiological outcomes. Together, our results demonstrate that within-patient evolution may be a general feature of both acute and chronic infections, and that gene gain/loss may be an important but under-appreciated feature of within-host evolution.

## Introduction

Cholera is an acute diarrheal infection that remains a serious health threat in countries with limited access to clean water [1]. *Vibrio cholerae* is the causative agent of the disease and is a natural inhabitant of aquatic ecosystems [2], with more than 200 serogroups identified to date on the basis of their somatic O antigens [3,4]. Most *V. cholerae* serogroups are not pathogenic; only isolates in serogroup O1 (consisting of two biotypes known as “classical” and “El Tor” and the serotypes Ogawa and Inaba) and O139 have been identified as agents of cholera epidemics and pandemics [1].

Whole genome sequencing and population genomics have the potential to improve our understanding of the epidemiology, etiology and evolution of bacterial infectious diseases [5]. For example, comparisons of whole-genome sequences of *V. cholerae* strains from across the world, over the course of a century, clarified the history of the current pandemic [6] and showed that this pandemic is the result of a single clonal expansion of one *V. cholerae* O1 El Tor ancestor, accompanied by horizontal gene transfer (HGT) events involving toxin and antibiotic resistance genes [7]. More recently, comparative genomics have been applied to answer epidemiological questions, proving the Asian origin of the strain causing the ongoing Haitian cholera outbreak, which began in 2010 [8-11]. Using whole genome sequencing and single nucleotide polymorphism (SNP) analysis, Azarian et al. [12] compared 60 clinical and environmental isolates collected in Haiti from 2010 to 2012. They found that the 2011 and 2012 strains rapidly diverged from the 2010 ancestral strain that initiated the outbreak, suggesting evolution driven by positive selection in a new environment [12].

Viral pathogens can evolve and diversify within infected patients, with serious consequences for disease outcome [13], and certain bacterial pathogens have recently been shown to diversify within patients as well [14]. However, evolutionary and epidemiological studies are usually conducted with just one bacterial isolate taken as representative of the infection, even though within-patient diversity is important to consider, for several reasons [15-18]. Within-host evolution may impact the longer-term evolution and transmission potential of pathogens, particularly if there are fitness tradeoffs between evolution within and between hosts. For example, a study of one cholera patient from Haiti showed that phage-resistant *V. cholerae* mutants rose to high frequency within the patient due to positive selection imposed by phage predation[19]. This study showed how strong selection can shape *V. cholerae* diversity within patients, but the prevalence and extent of *V. cholerae* genetic diversity within patients remains unclear, and whether intra-host evolution is generally driven by selection.

As for many other bacterial pathogens, the prevailing orthodoxy is that *V. cholerae* infections are clonal, and essentially devoid of within-host genetic diversity. Although within-host *V. cholerae* populations have not been studied extensively, additional evidence suggests that within-host diversity does indeed exist, at the level of phase variation in the O antigen, or in variable number tandem repeat (VNTR) loci [4,20]. This diversity could arise by within-host evolution, or be due to infection by different strains that diverged before entering the host. *V. cholerae* is genetically diverse in aquatic ecosystems [21] and co-infections from diverse environmental strains are possible [22]. *V. cholerae* infections are acute, lasting only a few days before the patient either recovers or dies [1]. Therefore, there is limited time for within-host evolution (including mutation, recombination, and selection) to occur. It is therefore expected that *V. cholerae* will experience less within-host evolution compared to more chronic bacterial infections with documented within-host evolution [23-28]. On the other hand, *V. cholerae* grows to large population sizes within the host (from 10^7^ to 10^9^ vibrios per gram of stool), dominating the gut microbiome [29,30]. If the effective population size within a host is large, many mutations are expected and natural selection will be efficient. However, *V. cholerae* likely experiences population bottlenecks upon infection and within the gut [31], which would reduce genetic diversity and reduce the efficiency of selection. In addition to point mutations, *V. cholerae* can undergo high rates of HGT [7,32,33], providing an additional potential source of within-host diversity. During an infection, *V. cholerae* could acquire genes from plasmids, phages, pathogenicity islands or genes from the gut microbiota, which appears to be a hotspot of HGT [34]. However, the extent of within-patient mutation, HGT, and natural selection are still poorly known for *V. cholerae*.

In this study, we quantified within-patient genetic diversity of *V. cholerae* infections. We characterized genomic diversity of *V. cholerae* within and between eight cholera patients, sequencing between eight and 20 isolate genomes per patient. We identified both intra-host single nucleotide variants (iSNVs) and gene gain/loss events within patients. As expected for an acute infection, few within-patient point mutations were detected, ranging from zero to three iSNVs per host. In contrast, we found a substantial amount of gene content variation: between five and 103 gene gains or losses within each patient. We infer that most diversity is due to within-host mutation rather than co-infection, and that HGT of mobile elements is more common than point mutations. In most patients, within-host evolution can be explained by neutral mutation, recombination and bottlenecks; however, one patient showed evidence for diversification driven by positive selection, resulting in phenotypic variation among intra-host *V. cholerae* isolates in their ability to form biofilms. Despite the relatively small numbers of mutations and HGT events within hosts, these events may have important evolutionary and phenotypic consequences for *V. cholerae* populations.

## Results

### Genomic characterization of within-patient isolates

We isolated and sequenced a total of 122 clinical strains recovered in 2013 from eight patients with culture-confirmed cholera. Five patients were from Dhaka, Bangladesh. Samples were collected during the same week at the end of 2013, and identified as patients B1, B2, B3, B4 and B5. The three other patients (H1, H2, H3) came from Artibonite, Haiti and presented with cholera, all during a three-day period in April 2013. For each patient, eight to 20 colonies were isolated from a single stool sample to assess within-patient genetic variation (Fig 1). Colonies were named with a C, followed by a number; for example, B1C1 corresponds to colony 1 from Bangladesh patient 1. All strains were identified as toxigenic *V. cholerae* O1 biotype El Tor, serotype Ogawa which was the prevailing serotype at each site during the entire study period. Each isolate was separately sequenced using Illumina technology to a minimum depth of 28X coverage of the MJ-1236 reference genome (mean coverage = 136X). We *de novo* assembled each genome, and also mapped reads to two closed, annotated *V. cholerae* reference genomes. After filtering for errors due to culture and sequencing (see Materials and Methods), we identified a total of 485 high quality single-nucleotide polymorphisms (hqSNPs) among our 122 isolates, distributed across the genome (S1 Fig), with a majority (471 hqSNPs) located in the branch between the Bangladeshi and the Haitian strains.

**Fig 1.**
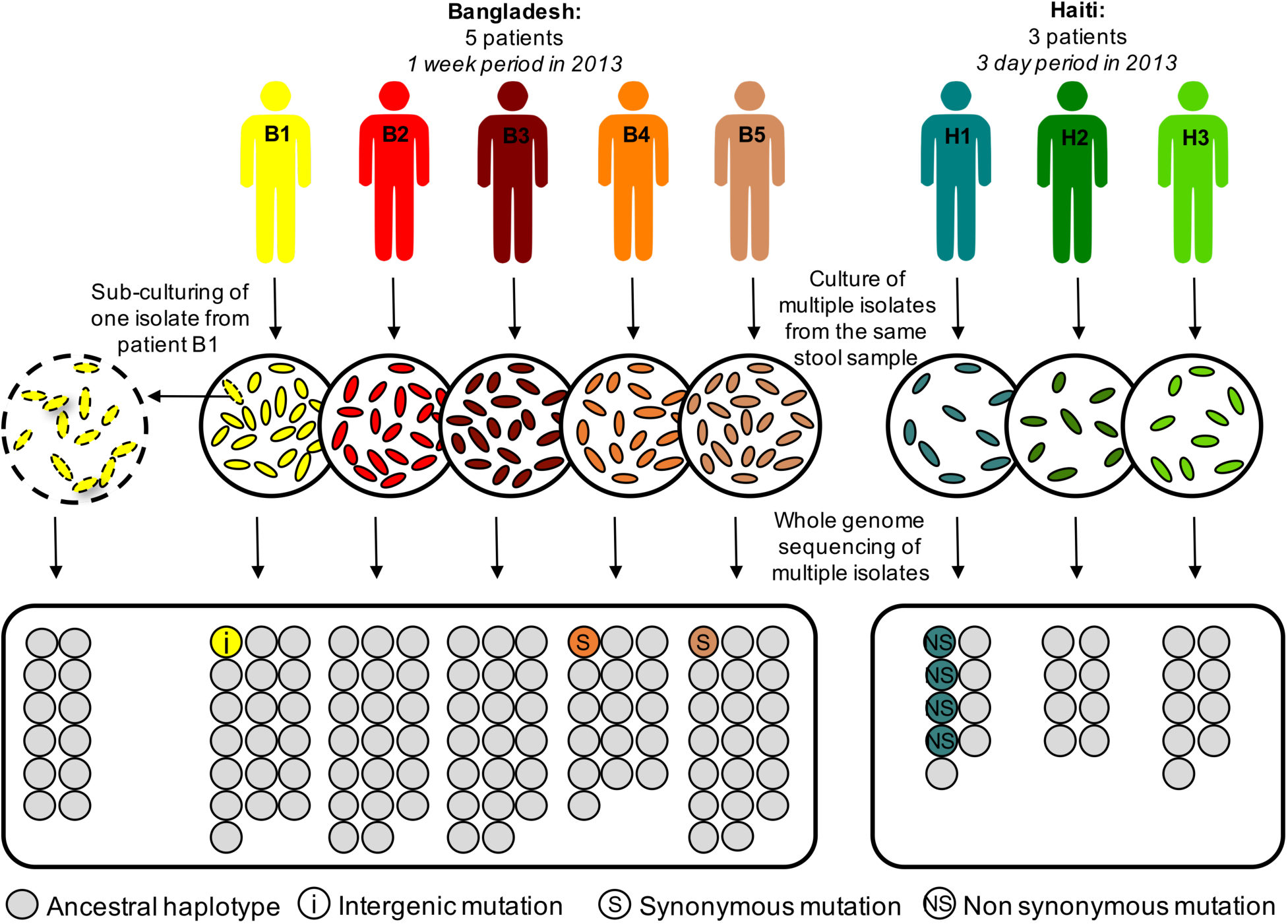
Culture and sequencing of *Vibrio cholerae* isolates from eight acutely infected patients. To study within-patient evolution, we cultured stool samples from five patients from Bangladesh (B1 to B5) and three patients from Haiti (H1 to H3), on selective media. We isolated between eight and 20 colonies from each patient and sequenced them separately. For patient B1, we performed a sub-culture of one isolate (dotted outline) and sequenced twelve of these new isolates, as a control for cultured-induced and sequencing artifacts. We independently called variants, compared them between isolates within each patient to identify the intra single nucleotide variants (iSNVs, colored circles) and determined whether they were intergenic (i), synonymous (S), or non-synonymous (NS) mutations.

### Within-patient single nucleotide variation

The set of hqSNPs included intra-host single nucleotide variants (iSNVs) in patients B1, B4, B5 and H1 (Fig 1). Isolates within patients B2, B3, H2 and H3 were isogenic, with no iSNVs detected using our quality filters. Patient B1 contained one intergenic iSNV, and patients B4 and B5 each contained a synonymous iSNV, each in a different gene (Table 1). Patient H1 contained three iSNVs, all of which were non-synonymous, and two of which occurred in the same gene, a sensor histidine kinase (Table 1). No small insertion/deletions (indels) were found within any patients.

**Table 1.** Nucleotide and amino acid changes identified in the *V. cholerae* core genome.

| Region | Type | Strains | Reference nucleotide | Alternative nucleotide | NS/S | Reference amino acid | Alternative amino acid | Gene annotation | Patient allele frequency |
| --- | --- | --- | --- | --- | --- | --- | --- | --- | --- |
| Bangladesh | iSNV | B1C19 | G | A | intergenic | - | - | - | 1/19 |
| Bangladesh | iSNV | B4C12 | G | A | S | D | D | RNA polymerase sigma factor | 1/17 |
| Bangladesh | iSNV | B5C11 | A | C | S | V | V | toxin co-regulated pilus biosynthesis protein F | 1/20 |
| Haiti | iSNV | H1C5 | C | A | NS | R | L | sensor histidine kinase | 1/9 |
| Haiti | iSNV | H1C4<br>H1C8 | C | T | NS | R | H | TetR family transcriptional regulator | 2/9 |
| Haiti | iSNV | H1C6 | C | A | NS | R | L | sensor histidine kinase | 1/9 |
| Bangladesh | Patient | B2 + B3 | A | G | intergenic | - | - | - | 20/20<br>20/20 |
| Bangladesh | Patient | B4 | G | A | NS | G | R | TatD family hydrolase | 17/17 |
| Bangladesh | Patient | B4 | G | T | NS | A | T | hypothetical protein | 17/17 |
| Bangladesh | Patient | B2 + B3 | C | T | intergenic | - | - | - | 20/20<br>20/20 |
| Bangladesh | Patient | B4 | G | A | intergenic | - | - | - | 17/17 |
| Bangladesh | Patient | B4 | C | T | NS | P | L | PTS system mannitol-specific EIICBA component | 17/17 |
| Bangladesh | Patient | B2 + B3 | C | T | NS | P | S | LacI family transcription regulator | 20/20<br>20/20 |
| Bangladesh | Patient | B1 + B5 | A | G | N | A | V | bifunctional purine biosynthesis protein PurH | 19/19<br>20/20 |

Mutations segregating within patients are denoted iSNVs; Mutations fixed between patients are
denoted 'Patient.' Patient allele frequency shows the allele frequency of the alternative (minor)
allele. NS=nonsynonymous; S=synonymous.

To ensure that the relatively small number of iSNVs were not due to mutation during strain isolation and culture [35] or sequencing errors, we sub-cultured and sequenced 12 colonies from one isolate (B1C1) as a control (Fig 1). Applying the same filters as for SNP discovery in our patients, we did not detect any iSNVs among control strains, nor did we detect any SNP differences between replicate libraries prepared and sequenced using different platforms (Materials and Methods). This suggests that the few iSNVs identified within patients are unlikely to be culture or sequencing artifacts.

Based on the number (0-3) and frequencies of iSNVs per host, we used measures of genetic diversity (θ_W_ and π) to estimate within-host effective population size (N_e_). We estimated that N_e_ within each patient ranged from 0 to 110 (S1 Table). Such a small N_e_ in a bacterial population is consistent with a recent population bottleneck, possibly during host colonization, or a recent selective sweep having purged most of the diversity within the population.

### Gene gain and loss within and between cholera patients

To characterize variation in gene content among the 122 sequenced isolates, we analyzed orthologous coding sequences from *de novo* assemblies. Of 3907 gene families identified, 3489 were defined as core (i.e. present in all genomes) while 401 (~10% of the total gene pool) were considered flexible, present in only a subset of genomes. Due to variation in sequencing coverage and nucleotide composition, genome assemblies may be incomplete, missing a subset of genes that are actually present [36,37]. To address this issue, we mapped reads back to the full gene catalogue, and considered a gene to be present when it was covered at 1X (Materials and Methods; S2 and S3 Fig). This filtering procedure revealed that, of the 401 genes initially classified as flexible, 252 were actually part of the core, leaving 155 *bona fide* flexible genes. Before filtering, we observed gene content variation among control isolates, but these false positives were removed using the 1X coverage filter. After filtering, between five and 103 flexible genes showed variation within patients (Table 2; Fig 2). Because gene content variation could be biased by different methods of library construction and sequencing, we sequenced 12 strains in duplicate using two different methods. We observed variation in the detection of the flexible genome for six of the duplicate sets (S4 Fig), likely because different library preparation methods are known to have different GC-content biases [38]. Using genomes from one method only (NEBNext/HiSeq), we identified between five and 67 variable genes per patient; using the other method (Nextera/MiSeq) we identified between zero and 62 genes (S2 Table). When a variable gene is not detected in a given method, we consider this a false negative. We conclude that methodological differences alone cannot explain the flexible genome variation within patients, and consider both methods combined for the remainder of the study.

**Fig 2.**
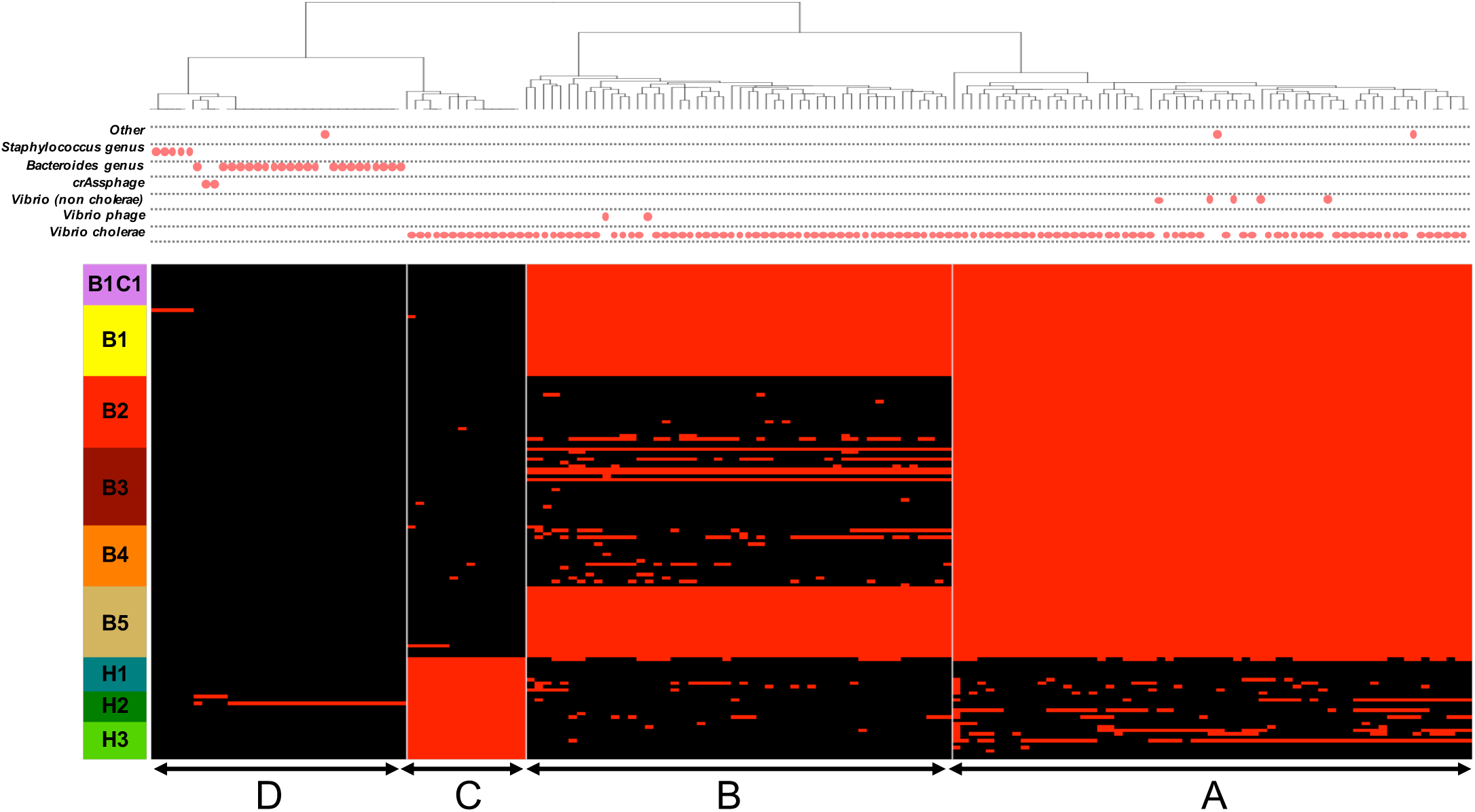
Presence/absence profile and taxonomic affiliation of gene families in the flexible genome. Red in the heatmap indicates gene presence; black indicates absence. Each column shows the presence/absence profile for a unique gene family. The heatmap is ordered by patient along the vertical axis. B1C1 is the control, subcultured from B1, and contains no flexible genome variation. The horizontal axis is ordered by hierarchical clustering, yielding four clusters: A, B, C and D. The taxonomic affiliation of each gene family (best BLAST hit) is indicated with red dots above the heatmap.

**Table 2.** Flexible gene content variation within and between patients.

| Patients | #genes fixed within<br>patients | #genes variable within<br>patients | #singletons |
| --- | --- | --- | --- |
| B1 | 111 | 11 | 5 |
| B2 | 61 | 35 | 0 |
| B3 | 61 | 51 | 0 |
| B4 | 61 | 49 | 0 |
| B5 | 111 | 5 | 0 |
| H1 | 14 | 68 | 0 |
| H2 | 14 | 103 | 25 |
| H3 | 14 | 63 | 0 |

Singletons are defined as genes only found in one strain, and are also counted as variable genes within patients. Genes fixed within patients are present in all isolates from a patient, but are absent in at least one other isolate in the study.

Clustering the 155 flexible genes by their presence/absence profile across patients revealed four distinct categories of genes (Fig 2). Category A consists of genes present in Bangladesh (part of the Bangladesh core genome), but showing a patchy distribution in Haiti. Category B genes are fixed (present in all isolates) in patients B1 and B5, and patchy in other patients. Category C genes are fixed in Haiti and nearly absent from Bangladesh. Category D genes tend to be rare, often singletons only observed in a single isolate within either patient B1 or H2.

Several flexible genes corresponded to known mobile genetic elements. Notably, category A contains 61 gene families (39% of the flexible genome), all located on one single contig corresponding to the SXT Integrative Conjugative Element (ICE). Category B encompassed 49 gene families matching Kappa phage proteins. These putative phage genes were clustered together on large contigs of chromosome 1, and were fixed in patients B1 and B5 but variable among other patients, some of which contained complete phage sequences (patients B2 and B3). Category C contained 15 genes, including some present in the ICE, which mapped to at least five different contigs (depending on which isolate’s assembly was considered), suggesting multiple gain/loss events or frequent rearrangements.

Over half of the flexible genome (80 genes) was annotated as hypothetical proteins, compared to the core genome that contained less than 3% hypotheticals. The flexible genome also contained ten transposases (6.5% of the flexible genome, compared to 1.4% of the core) and eight genes involved plasmid and viral replication, all potential mechanisms of HGT[39]. A complete list and annotation of flexible genes is given in Supplementary Table 5.

Variation in the flexible gene pool could arise from gene deletion, duplication, or horizontal gene transfer (HGT). To determine the extent of HGT across species boundaries, we identified the taxonomic affiliation of each flexible gene according to its best BLAST hit in GenBank. While 117 of the 155 of flexible genes were assigned to *V. cholerae*, the other 38 genes were assigned to other members of the family *Vibrionaceae* or even distantly related species of *Bacteroides* or *Staphylococcus* (Fig 2), suggesting HGT from these donors to *V. cholerae* in the gut. For example, a group of 20 genes present in isolate H2C3 (but absent in other isolates from patient H2) matched a plasmid previously identified in *Bacteroides* (Fig 2). These 20 genes of putative *Bacteroides* plasmid origin are among 25 singletons, present in only one isolate of patient H2. Similarly, the 5 singletons in patient B1 (Table 2) are all of putative *Staphylococcus* origin. As no other genes were assigned to either *Staphylococcus* or *Bacteroides*, it appears these putative HGT events are not due to contamination from these taxa. This suggests that cross-species HGT events are relatively rare and recent events, which may never achieve high frequency in the *V. cholerae* population, either within or across hosts. Together, these results suggest that most within-patient variation in gene content is due to gene flow, deletion or duplication within the *V. cholerae* population, with rare but detectable HGT from other species of bacteria, phages and plasmids in the gut microbiota.

### *V. cholerae* evolution on different time scales

In order to place within-host variation in the context of longer-term *V. cholerae* evolution, and to distinguish within-patient mutation from coinfection events, we built a phylogeny of the 122 isolates (all from 2013) as well as 21 additional isolates obtained from acute cholera patients sampled in Bangladesh from 2011 to 2013 (Fig 3). These 21 clinical isolates were processed with the same methods as the within-host isolates, from the culture to the hqSNP discovery (Materials and Methods). With a total of 35 unique genotypes (one to four per patient), we computed a Bayesian phylogeny, estimated divergence times, and evolutionary rates using BEAST 1.8.3 [40]. We used an alignment of all the concatenated hqSNPs except those found in the ICE, a highly variable 100 kb region (S1 Fig) that undergoes frequent HGT and thus has a separate evolutionary history from the rest of the genome [6].

**Fig 3.**
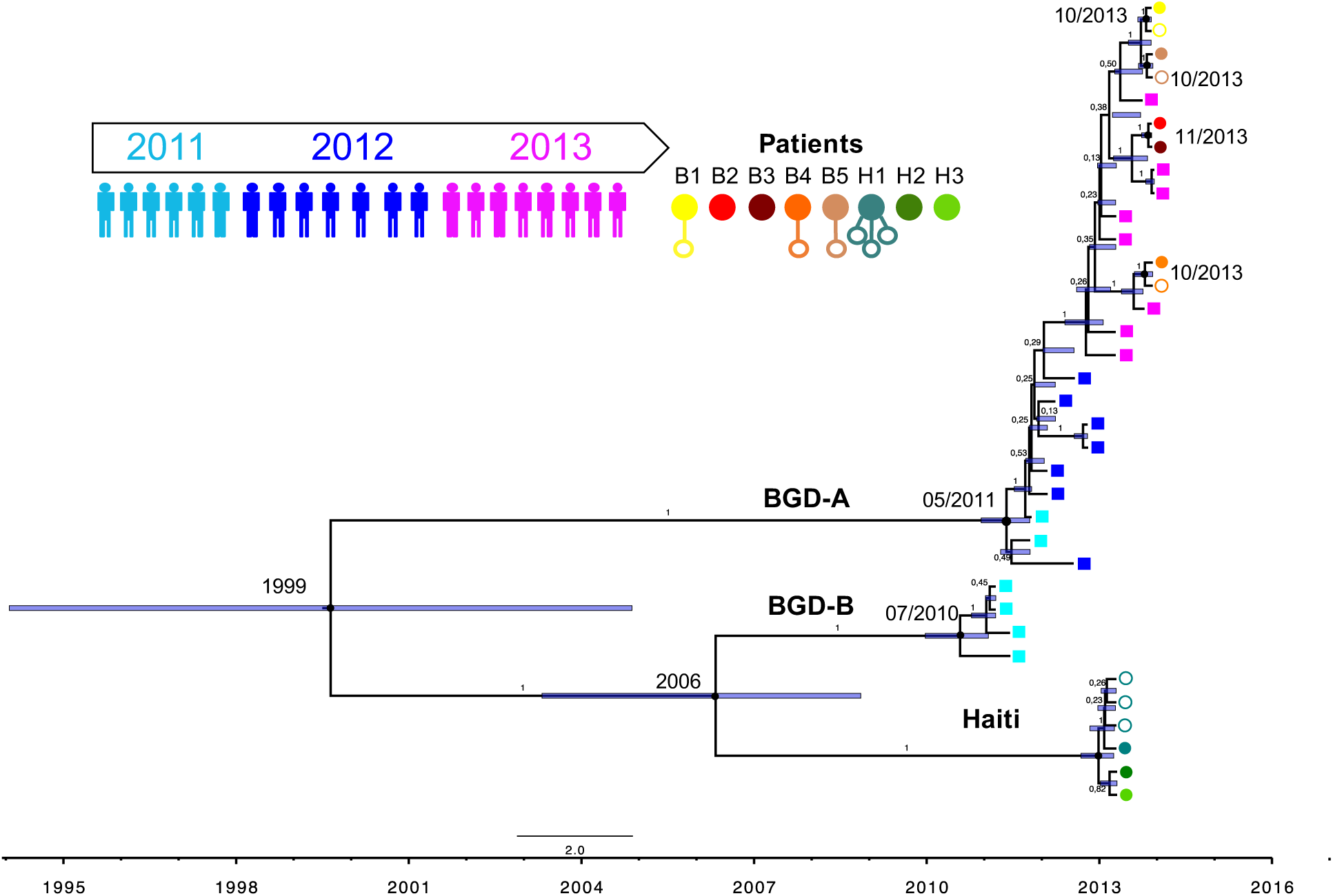
Bayesian phylogenic tree of 35 *V. cholerae* genotypes sampled over three years in Bangladesh and Haiti. The maximum clade credibility tree represents the genealogy of sequences in the study, reconstructed from concatenated hqSNPs, using BEAST. Colored squares (shades of blue and purple) represent the time-course isolates collected from Bangladeshi patients from March 2011 to December 2013 (one isolate per patient). Patients for whom we measured intra-host variation (B1-B5 and H1-H3) are shown as circles. Filled circles indicate the putative ancestral genotype, and empty circles indicate putatively derived iSNVs. The median node age and divergence date in months and years are indicated at the nodes. The blue bars represent the 95% HPD intervals for divergence time estimates, and posterior probabilities are represented on the branches.

To test the degree of temporal structure in the phylogeny, we performed a root-to-tip linear regression (R^2^=0.65, p<0.0001; S5 Fig), which suggested that our dataset is sufficiently clock-like to robustly estimate an evolutionary rate. The Bayesian tree was calibrated with sampling dates of the isolates ranging from March 2011 to December 2013 (Fig 3). We compared different molecular clock and demographic models (Materials and Methods), and found that the strict molecular clock and the Bayesian skyline plot models provided the best fit (S3 Table), in accordance with previous studies of *V. cholerae* [12,41]. Assuming a constant evolutionary rate across lineages, we estimated the evolutionary rate at 7.94 × 10^−7^ hqSNP site^-1^ year^-1^ (95% highest posterior density [HPD], 4.89 × 10^−7^ to at 1.14 × 10^−6^) or approximately 3.3 hqSNP year^-1^ in the core genome (95% HPD), consistent with previous estimates [6].

We then estimated the ages of the most recent common ancestors (MRCAs) of the phylogenetic sub-lineages and clusters (S4 Table). Notably, four of the six strains isolated in 2011 from Bangladesh were found to be closer to the Haitian strains than to all the other Bangladeshi strains (Fig 3). We called this sub-lineage BDG-B, and estimated the age of the MRCA of these four strains and the Haitian isolates in September 2005 (95% HPD, June 2002 to July 2008), which pre-dates the introduction of pathogenic *V. cholerae* in Haiti, and is consistent with its Asian origin [10,11]. The time of the MRCA of the strains isolated from the three Haitian patients was estimated at December 2012 (95% HPD, August 2012 to March 2013).

Based on the phylogeny (Fig 3), we sought to distinguish between scenarios of within-patient mutation or coinfection as causes of within-patient diversity. It is clear that isolates from the same patient always grouped together, and were never polyphyletic. This observation is consistent with each patient being colonized by a single clone, which subsequently diversified by mutation within the patient. The diversity between the eight patients (7 SNPs) was greater than the diversity within patients (0-3 iSNVs), which would be unlikely if strains were sampled by patients at random from an environmental gene pool (Table 1). If within-patient diversity was due to coinfection of the same patient by multiple different strains, we would expect these strains to share a MRCA before the date of infection, and certainly before the date stool was sampled. However, the MRCA of isolates from a single patient always overlapped with the date of sampling, suggesting that within-patient diversity is more likely due to within-patient mutation than to coinfection.

### Signatures of natural selection on within-patient variants

We next asked how regimes of natural selection on *V. cholerae* varied over time, within and between patients. We used the nonsynonymous (NS) to synonymous (S) substitution ratio to measure the strength of selection at the protein level. Over three years of evolution, we could not reject a neutral evolutionary model and found no evidence for variation in the NS:S ratio over time, considering only SNPs fixed between patients (Supplementary Text). Another possibility is that selection acts over shorter evolutionary scales, by shaping intra-host diversity during acute infection. Under this scenario, we would expect NS:S ratios to differ significantly within and between hosts. For example, higher NS:S within than between hosts could be due to positive or balancing selection on NS mutations within hosts, or due to more efficient purifying selection (against deleterious NS mutations) between hosts. To test for such deviations from neutral evolution, we applied the McDonald-Kreitman test [42] to the eight hosts surveyed for within-host genetic variation (five from Bangladesh and three from Haiti). Despite the overall low number of SNPs and iSNVs observed, we found a significant excess of NS mutations between Bangladeshi patients (Fisher’s exact test, Odds Ratio > 12, p<0.05; Table 3), suggesting positive selection for the fixation of NS mutations between patients, or purifying selection against NS mutations within patients. In contrast, all three iSNVs observed in Haiti were NS, suggesting positive, balancing, or relaxed purifying selection within patients, although not statistically significant (Fisher’s exact test, Odds Ratio < 0.32, p=1; Table 2).

**Table 3.**
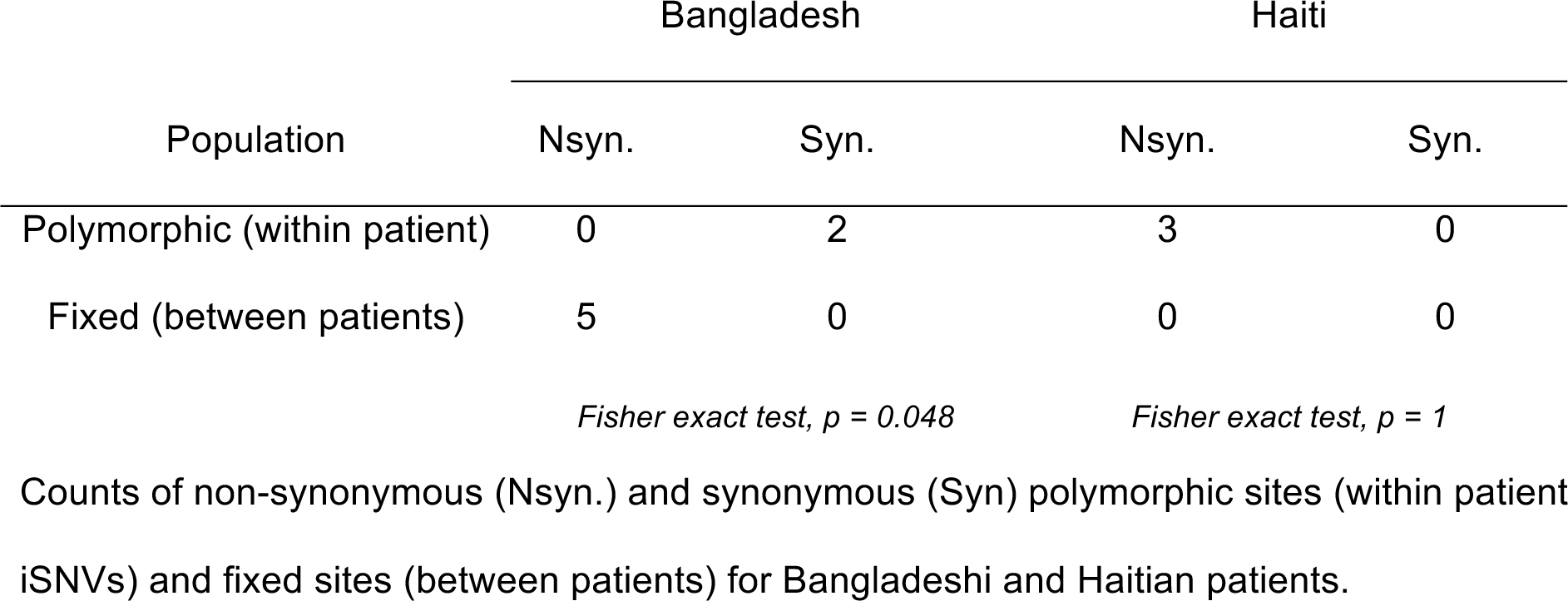
McDonald-Kreitman test for differential selection within and between patients.

The three NS iSNVs observed in Haiti all occurred within a single patient, possibly driven by selective pressures specific to this patient (Table 1). To test whether this pattern of iSNVs was likely to have occurred at random or due to patient-specific selection, we performed permutations of iSNVs among hosts and estimated expected iSNVs frequencies (*F*) and number of NS iSNVs per host and region. We found a significant excess of iSNVs in Haiti (*F*_HTI_ = 0.15; p<0.05; 10,000 permutations) and in patient H1 (*F*_H1_ = 0.44; p<0.01), but not in Bangladesh (*F*_BGD_ = 0.03; p>0.05) nor in any other patients (*F* = 0-0.05; p>0.05). All iSNVs identified in Haiti and in patient H1 were non-synonymous, which was significantly higher than expected by chance (p<0.01 and p<0.001, respectively; Fig 4). These results show that patient H1 has a significant excess of NS iSNVs compared to other patients. This suggests positive or balancing selection on NS iSNVs within patient H1, or relaxed purifying selection in H1 compared to other patients.

**Fig 4.**
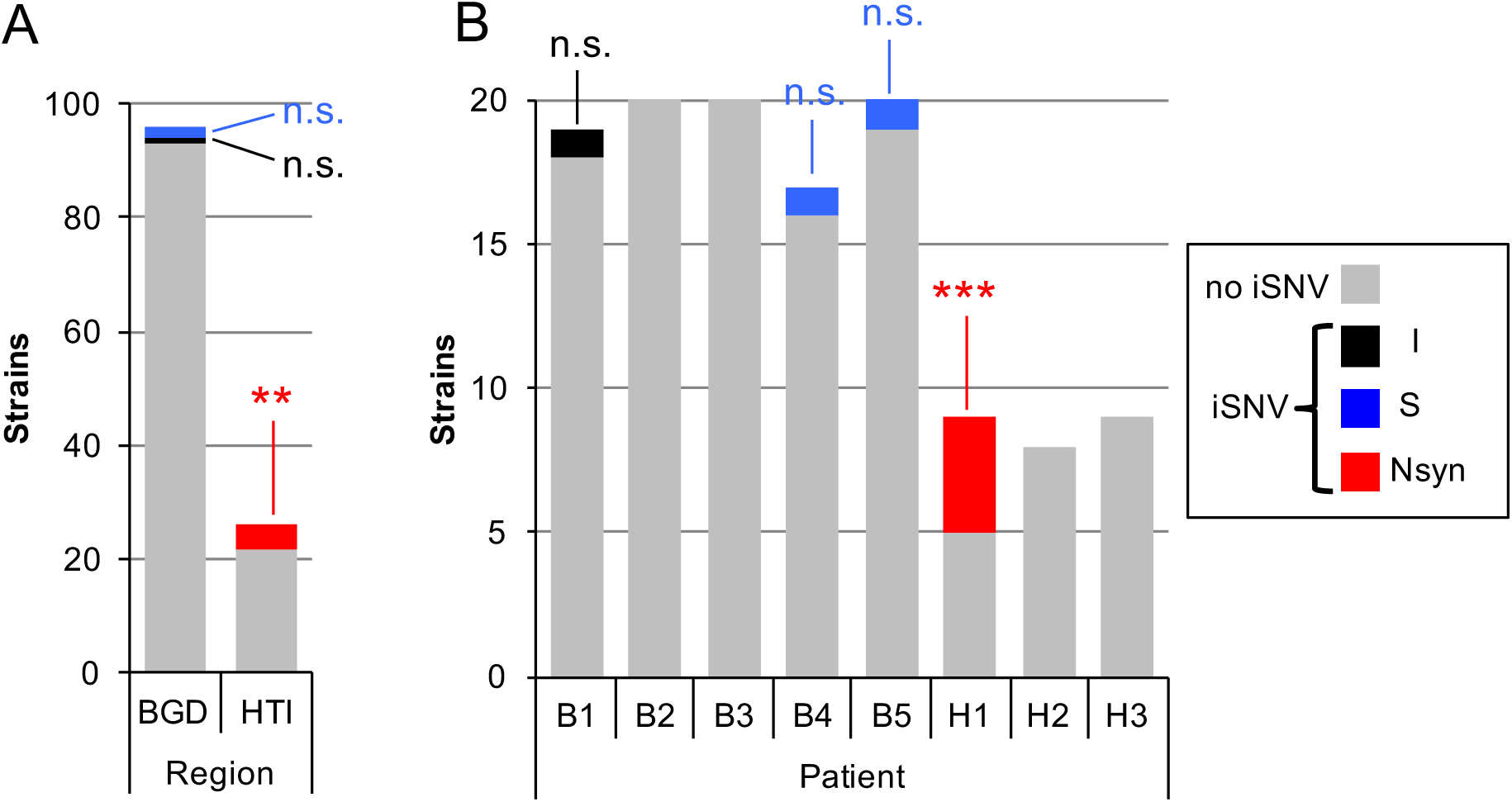
Significant excess of non-synonymous iSNVs in patient H1. (A) Distribution of 122 *V. cholerae* strains containing different categories of iSNVs (I: intergenic; S: synonymous; Nsyn: non-synonymous) or no detectable iSNVs, according to geographic region (BGD: Bangladesh; HTI: Haiti). Patients from Haiti have a significant excess of Nsyn iSNVs (red; *p*≤0.01; 10,000 random permutations of strains among regions). (B) Distribution of 122 *V. cholerae* strains containing different iSNVs, or no detectable iSNVs, by patient. Patient H1 has a significant excess of strains with Nsyn iSNVs (*p*≤0.001; 10,000 random permutations of strains among patients).

The three NS iSNVs in patient H1 occurred in two genes. The first gene, containing one iSNV, encodes a member of the tetracycline resistance (Tet^R^) family of transcriptional regulators (NCBI: ACQ60802.1), known to be involved in the transcriptional control of multidrug efflux pumps and other pathways like quorum-sensing circuits or pathogenicity[41,43]. The other two non-synonymous mutations in patient H1 were located in the same gene (NCBI: ACQ61177), a sensor histidine kinase (HK) called *vprB*, which is required for resistance to the antimicrobial peptide polymyxin B [44]. Each of these two iSNVs occurs at a different site in the gene, each in a different isolate. Based on the fact that the major allele at each of these iSNV sites was present in both reference genomes (MJ1236 and 2010EL-1786), we inferred that the minor alleles (both at frequency 1/9 in patient H1; Table 1) were derived, presumably due to within-patient mutation. A comparison of the *vprB* (ACQ61177) protein sequence with its 465 closest orthologs revealed that the NS iSNVs identified in this gene modify peptides that are otherwise highly conserved across *Vibrio* species (S7 Fig), suggesting that these mutations may affect protein function.

### Within-patient variants affect biofilm formation

To identify possible phenotypes conferred by these within-patient variations, we performed three sets of experiments on the strains from patient H1 harboring the three NS point mutations, and the strain for which the *Bacteroides* plasmid was detected in patient H2. First, to test whether intra-host variants had any effect on *V. cholerae* growth in liquid medium, we measured the growth rates of strains with derived iSNV alleles compared to one isogenic strain (with inferred ancestral alleles and no variation in the flexible genome) from the same patient, and found no significant difference in the growth rates between strains. Second, we tested for resistance to the antibiotic polymyxin B, as loss of *vprB* function has been previously associated with increased susceptibility to polymyxin B [44], and again found no difference between strains (S8 Fig). Third, we tested for biofilm formation, as it has been previously suggested that biofilm formation inside the host can impair intestinal colonization [45,46], and it is known that HKs in certain two-component systems can affect biofilm formation (48,49). As a biofilm-negative control, we used RBM, a biofilm knock-out strain of *V. cholerae* [47], sterile LB, or saline.

For patient H1, we did not detect any significant difference in biofilm formation between strains with the ancestral iSNV allele (in isolate H1C1) and the derived allele in the transcriptional regulator gene (isolate H1C4). However, we found that H1C5 and H1C6, the two isolates with derived alleles in the HK gene *vprB*, produced significantly less biofilm than the other strains from the same patient (Fig 5A). Based on our hqSNP calls and flexible gene analysis, the genome of H1C5 was identical to H1C1, with the exception of the iSNV in the HK gene. Therefore, the difference in biofilm phenotypes is attributable to this iSNV. However, H1C6 differed from H1C1 by an iSNV in the HK gene, and also by the presence/absence of genes in the flexible genome (S4 Fig). However, this gene content variation did not measurably affect biofilm formation (Fig 5A).

**Fig 5.**
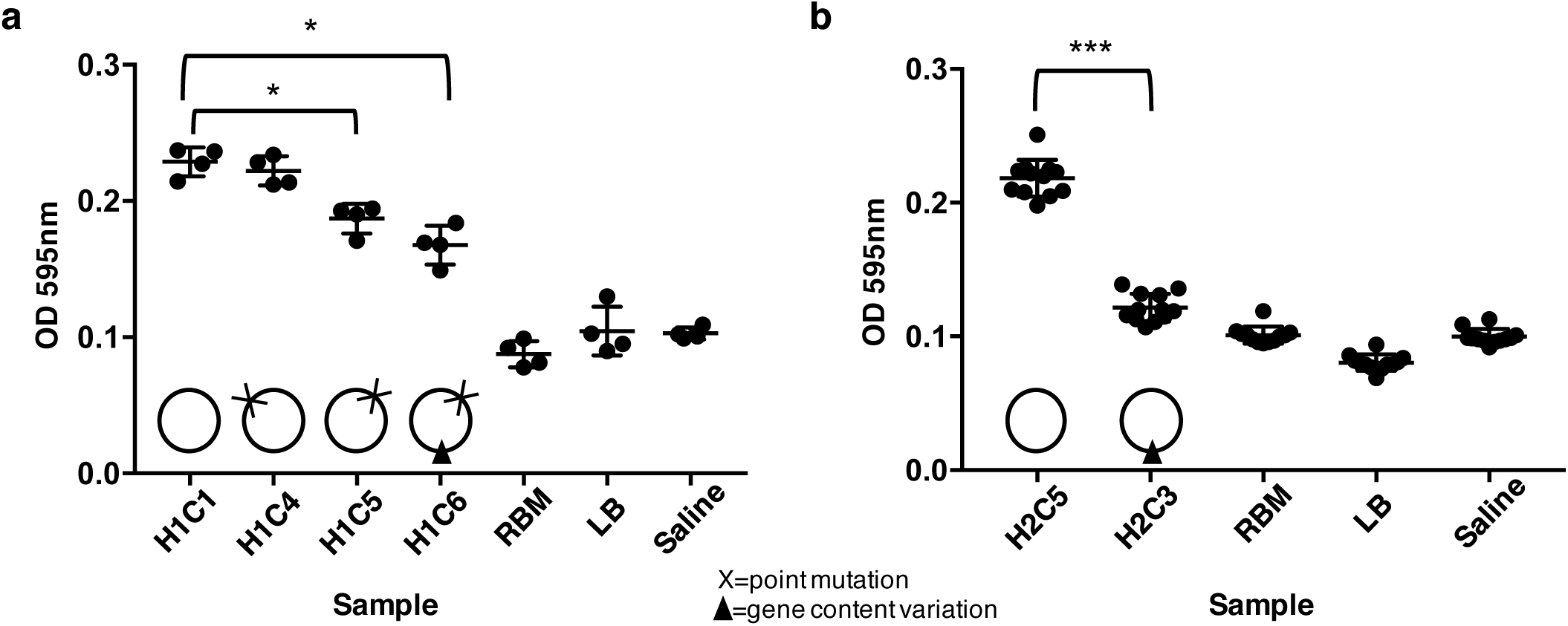
Biofilm formation of strains from patients H1 and H2. Optical density was measured for four to 12 replicates of each strain, after 48h of growth at 30°C. Statistical comparisons were made using a non-parametric Mann-Whitney test (^∗^*P* < 0.05, ^∗∗∗^*P* < 0.0001). Circles represent genomes with either variation in gene content (dark triangle) or iSNV variation (cross), **(a)** Strains from patient H1. Isolate H1C1 represents the ancestral genotype, H1C4 has a nonsynonymous mutation in a transcriptional regulator gene, and H1C5 and H1C6 have different nonsynonymous mutations in the same gene, the histidine kinase gene, **(b)** Strains from patient H2. Isolate H2C5 represents the ancestral genotype, with no variation in the gene content, and H2C3 harbors a plasmid. RBM is a biofilm knockout strain, and LB and saline are negative controls.

In the case of patient H2, we showed that the presence of a plasmid of putative *Bacteroides* origin strongly affects biofilm production. Specifically, the plasmid-containing isolate (H2C3) produces approximately two-fold less biofilm than an isogenic control from the same patient (H2C5). H2C3 biofilm formation was indistinguishable from negative controls (Fig 5B). Together, these results show that both point mutations and plasmids segregating within patients can affect biofilm formation.

## Discussion

In this study, we surveyed the genetic diversity of *Vibrio cholerae* within infected patients. Using whole-genome sequencing, we analyzed 122 clinical isolates from eight cholera patients from Bangladesh and Haiti, and demonstrated that overall levels of within-patient variation are low for *V. cholerae* populations compared to more chronic bacterial pathogens, which routinely harbor more than 20 iSNVs per patient [24,25,26,27,28,48]. Even if rare, point mutations may be under selection within hosts, with phenotypic consequences. For example, we showed that intra-host mutations in a sensor histidine kinase gene reduced biofilm formation. In addition to point mutations, HGT plays a major role in *Vibrio cholerae* evolution and may represent the major source of genetic diversity, not only in the aquatic environment, but also in the human host – and with large effects on phenotypes like biofilm formation. Specifically, different mutations in a sensor histidine kinase and the acquisition of a plasmid both reduced the ability of *V. cholerae* to form biofilms, which could be advantageous during host colonization[44,45].

### Gene content variation within patients and its functional consequences

While HGT is already well-characterized on longer epidemiological time scales, we show that it also occurs within individual patients. *V. cholerae* is known to undergo HGT via transformation [49], transduction [50] and conjugation [51,52]. HGT contributes substantially to drug resistance, pathogenicity and adaptation to different environments, via the acquisition of genomic islands, phages, transposons, ICEs and plasmids [7,53]. Our characterization of the flexible genome within patients used read mapping to confirm gene absences, reducing false positive inference of gene content variation (S2 and S3 Fig). We detected between five and 103 genes that varied in presence/absence within patients (Fig 2; Table 2). Each gene does not necessarily represent an independent gain/loss event; for example, the ICE contained 61 genes on a single contig, likely a single gain/loss event. Even under the conservative assumption that all gene content variation represents a single gain/loss event per patient, this still indicates at least one event per patient.

Some of the putative HGT events could have consequences for *V. cholerae* survival and virulence within the host. For instance, the group of 20 genes acquired by one *V. cholerae* isolate within patient H2, likely via a plasmid of *Bacteroides* origin, is associated with a two-fold reduction in biofilm formation (Fig 5). Among these 20 genes, we identified an antibiotic resistance gene, a haloacid dehalogenase protein that could impact pathogenicity [54], and a FtsY recognition signal protein that was shown to increase virulence in *Streptococcus* [55], among others (S4 Fig). Among the genes likely acquired from a *Staphylococcus* donor in isolate B1C2, three could potentially be involved in modulation of virulence (S4 Fig, S6 Table). First, a GNAT family acetyltransferase could promote virulence or increase antibiotic resistance [56,57]. Second, a putative phosphoenolpyruvate phosphotransferase has been demonstrated to play a role in biofilm formation in *Vibrio* [58,59,60]. Finally, the KdpC gene (a potassium-transporting ATPase) has been shown to modulate virulence in *Mycobacterium paratuberculosis* [61].

Not all gene gain/losses are due to HGT. Many can be explained by gene deletions, such as phage excision events. Deletions could explain much of the variation among genes in categories A and B (Fig 2), respectively corresponding to the ICE and Kappa phage. These elements are known to vary among *V. cholerae* strains, but here we document likely excision events during human infection. Genes in category D tend to be singletons, present in just a single isolate, and with taxonomic affiliations well beyond *V. cholerae*, including *Bacteroides*, *Staphylococcus*, and crAssphage (Fig 2). Category D genes are most easily explained by cross-species HGT, as previously documented by Folster and colleagues, who identified a Haitian isolate having gained multidrug resistance through transfer of a plasmid from a species of *Enterobacteriaceae* [51]. Taken together, these results are consistent with the human gut being a hotspot of HGT [62], sometimes involving pathogens like *V. cholerae*.

### Regimes of natural selection inferred from within-patient point mutations

The low levels of polymorphism (0-3 iSNVs per patient) we observed within cholera infections could be easily confounded with sequencing errors or mutations occurring during culture rather than within patients. Therefore, we developed filters for calling SNPs and gene gain/loss events that yielded zero variation among control colonies, suggesting low rates of false-positive variant calls and increasing confidence that the six total iSNVs (Fig 1; Table 1) did indeed vary within patients.

Point mutations detected within cholera patients could be the result of *de novo* mutations occurring within the patient, or a consequence of a co-infection from different strains that had diverged previous to the infection. Although we cannot formally exclude coinfections, our results are more consistent with *de novo* mutation within hosts. Isolates from the same patient were grouped together on the phylogeny (Fig 3), suggesting a clonal ancestor. This result is consistent with previous findings that cholera outbreaks are highly clonal [63].

Cycles of transmission from host to host, or from aquatic environment to host, can induce population bottlenecks, reducing the effective population size. Based on comparisons of distantly related genomes, N_e_ of *V. cholerae* has been estimated to be 4.78×10^8^ [64]. This relatively large N_e_ reflects the high genetic diversity present in the aquatic environment, and over long evolutionary time sales. However, during transmission and intestinal colonization, the size of the *V. cholerae* population experiences drastic bottlenecks that could temporarily reduce N_e_. Abel and colleagues showed that *V. cholerae* population sizes in rabbit models of infection ranged from 10^5^ during the early phases of colonization to ~10^2^ at the late phases of infection [31]. Our estimates of N_e_ based on iSNVs give values of 0 to ~10^2^, consistent with population bottlenecks or selective sweeps purging diversity (S1 Table).

In addition to the small within-patient N_e_, we found that the distribution of mutations, physically along the *V. cholerae* genome, and temporally along the phylogeny, could generally be explained by random neutral simulations (Supplementary text). However, we identified an excess of nonsynonymous mutations in one Haitian patient (H1), suggesting positive or diversifying selection on *V. cholerae* within this patient (Fig 4). Two of these mutations affected the same protein, a sensor protein-histidine kinase. This sensor protein histidine kinase (HK) is part of a two-component system known as VprAB, which has been shown to mediate glycine fixation in the lipid A domain of lipopolysaccharide molecules, which is necessary for resistance to the antimicrobial peptide polymyxin B [44]. Another two-component system, CarRS, is known to confer polymyxin B resistance, but also to negatively regulate biofilm formation [65,66]. Here, we found that derived alleles (presumed *de novo* mutations within the host) in the HK VprB did not appear to affect polymyxin B resistance (S8 Fig), but did reduce the ability of *V. cholerae* to form biofilms (Fig 5). It has been previously suggested that biofilm formation may be beneficial for survival in the aquatic environment, but detrimental to survival or colonization of mammalian hosts [45,46]. Therefore, the derived iSNV alleles may have been selected for reduced biofilm formation.

Our study provides an opportunity to compare *V. cholerae* evolutionary dynamics between Bangladesh, where cholera has been endemic for hundreds or thousands of years [67], and Haiti, where it was introduced in 2010. Our results suggest that selective pressures on *V. cholerae* may differ between Haiti and Bangladesh, as previously proposed [12]. In Bangladesh, we observed an excess of nonsynonymous (NS) mutations between patients (Table 3), suggest positive selection on protein sequences between patients, or efficient purifying selection purging nonsynonymous mutations within patients. In contrast to Bangladesh, where zero NS iSNVs were observed, we observed three NS iSNVs in Haiti, all within the same patient. Such differences between Haiti and Bangladesh need confirmation in a larger sample, and if confirmed could have different explanations. For example, if Haitian patients are less likely than Bangladeshis to have had prior exposure and immunity to cholera, perhaps cholera infections could last longer, or support larger *V. cholerae* population sizes within patients in Haiti, allowing more efficient positive selection within patients. Further sequencing of intra-host *V. cholerae* genomes, ideally in combination with clinical data on infection durations and outcomes, will be needed to test this hypothesis.

### Conclusion

In summary, we have shown that small but measurable genomic changes occur in the *V. cholerae* genome during human infections. Changes in flexible gene content appear to accumulate more quickly than point mutations, although point mutations may be targets of natural selection. Both gene content variation and point mutations can have consequences for the phenotypes of within-patient *V. cholerae* populations, including clinically-and environmentally relevant traits like biofilm formation. Future studies will be necessary to determine the role of intra-host diversity in the evolution of antibiotic resistance, host adaptation, and the severity of disease in infected patients.

## Materials and methods

### Enrollment

To study cholera within-host diversity, stool samples were collected from five patients (B1 to B5) from Dhaka, Bangladesh, and three patients (H1 to H3) from Artibonite, Haiti. Between eight and 20 *V. cholerae* colonies were isolated from each patient, as described below. Patients in Bangladesh were enrolled at the icddr,b (International Center for Diarrheal Disease Research, Bangladesh) Dhaka Hospital. The icddr,b cares for more than 120,000 patients annually including approximately 20,000 with cholera. Patients presenting during 2013 with acute watery diarrhea were eligible for inclusion in this study if stool cultures were positive for *V. cholerae* as the only pathogen, if they were between 2 and 60 years of age, resided in or around Dhaka, and were without major comorbid conditions. In Haiti, samples were collected from patients presenting to St. Marc’s Hospital, Arbonite, Haiti, with acute watery diarrhea in April 2013.

In addition to these eight patients, we included 21 “Time Course” patients (TC01 to TC21) from a surveillance program conducted by the icddr,b, between 2011 and 2013. For the Time Course samples, only one isolate was sequenced per patient.

### Sample Processing

In Bangladesh, diarrheal samples were examined by dark field microscopy and if positive for *V. cholerae* on presentation, then stool was cultured overnight. Samples with visible *V. cholerae* growth were serologically confirmed by slide agglutination with specific monoclonal antibodies for Ogawa or Inaba serotypes [68]. Confirmed cholera stool was stored in glycerol at −80°C and shipped to Massachusetts General Hospital, Boston. In Haiti, fresh stool from suspected cholera patients was stored in glycerol at −80°C and shipped from St. Marc’s Hospital to Massachusetts General Hospital, Boston. Stool from both Haiti and Bangladesh was stored at MGH at −80°C and then streaked directly onto thiosulfate-citrate-bile salts-sucrose agar (TCBS), a medium selective for *V. cholerae*. After overnight incubation, twenty well-separated colonies were inoculated into 5 mL Luria-Bertani broth and grown at 37°C overnight. For each colony, 1mL of broth culture was stored at −80°C with 30% glycerol until DNA extraction. For patient 1, one of the colonies was re-streaked on a new TCBS plate and 12 colonies were selected as a control for culture-induced artifacts and sequencing errors.

### DNA extraction and whole genome sequencing

Bacterial stocks made from a single colony were grown in 1.5mL LB media with agitation at 37°C for 12 hours. Genomic DNA was extracted for each isolate using the Qiagen DNeasy Blood^®^ and Tissue kit, using 1.5mL bacteria grown up in LB media. In order to obtain pure gDNA templates, an RNase treatment was followed by a purification with the MoBio PowerClean^®^ Pro DNA Clean-Up Kit.

From 122 isolates retrieved from the eight patients, 66 genomic libraries were constructed using the Nextera DNA library kit, according to the manufacturer’s protocol (Illumina) and were sequenced with the 250-bp paired-end v2 kit on the Illumina MiSeq. The remaining 56 libraries were prepared using the NEBNext Ultra II DNA library prep kit and sequenced on the Illumina HiSeq 2500 (paired-end 125 bp) at the Genome Québec sequencing platform (McGill University). Twelve isolates were sequenced in replicate using both methods. For details about isolates, sequencing and assembly see S5 Table.

### Genome assembly

To exclude low-quality sequences, we filtered raw reads with Trimmomatic [69]. The 15 first bases of each read were trimmed and reads containing at least one base with a quality score of <30 were removed. *De novo* assembly was then performed for each strain using IDBA_ud v1.1.1[70].

Filtered reads were also mapped with Bowtie2 v2.2.5[71] to a total of 11 references: two annotated reference genomes, one from Haiti (2010EL-1786, accession no. NC_016445.1 and NC_016446.1), one from Bangladesh (MJ1236, accession number NC_012667 and NC_012668), and nine assembled genomes (one from each patient and one from the sub-cultured control colonies: B1C1-06, B1C7, B2C12, B3C12, B4C5, B5C10, H1C5, H2C3, H3C2). PCR duplicates were removed from mapping using the MarkDuplicates function of PICARD TOOLS v1.130 (https://broadinstitute.github.io/picard/) and the SAMtools view utility [72], and realignment around indels was performed using the Genome Analysis Toolkit v3.1.1, with default settings. To facilitate the identification of homologous regions among the eleven reference genomes, MJ1236 and the nine *de novo* assembled genomes were aligned against the 2010EL-1736 genome using the “move contig” option in Mauve v. 2.4.0 [73], with default parameters.

### Variant calling and annotation

We used SAMtools v1.3 and BCFtools v1.1 to call SNPs and indels from mapping, requiring a minimum mapping quality of 30 and a minimum base quality of 20. Resulting SNPs and indels were then filtered by quality score (<20), depth of coverage (<10) and FQ scores (<0, lower values indicate agreement between reads) with VCFlib [74]. For each strain, we also filtered variable positions that were not retrieved in all the mappings against all 11 references, by performing reciprocal BLAST of the 50 nucleotides upstream and downstream of each variable position and comparing them. Only matches with >95% identity were kept and multiple matches were excluded as possible duplications and repeated elements. After applying these filters, we compared the genomes of the 12 replicate clones from the strain B1C1, to control for possible SNPs due to mutations during culture, or sequencing errors. We also removed positions that were called as variable when reads from one isolate were mapped to the assembly from the same isolate. We considered these SNPs as potential sequencing, mapping or assembly errors. Using these filters, we generated a list of high-quality SNPs (hqSNPs). From this list, we identified intra-host single nucleotide variants (iSNVs) as SNPs that were polymorphic among isolates from the same patient. No iSNVs were identified among the control colonies (the 12 replicate clones subcultured from B1C1).

Annotations were available for the MJ1236 reference genome and were retrieved from GenBank files (Chromosome 1: CP001485.1; chromosome 2: CP001486.1). Variants from the core genome were classified in three categories: intergenic (INT) when falling outside of a coding region, synonymous (S) when affecting the nucleotide sequence of at least one gene but not its amino-acid sequence; or non-synonymous (NS) when affecting the amino acid sequence of at least one gene.

### Estimation of effective population sizes (N_e_)

To estimate effective population size within each patient, we used the formula N_e_ = θ/2 μ, where N_e_ is the effective population size, θ is a measure of genetic diversity and μ is the mutation rate [75]. We assumed a mutation rate of μ=1/300 per genome per generation [76]. We report the estimated N_e_ for each patient in S1 Table, using both Waterson’s estimator (θ_W_ or S) and Tajima’s estimator (θ_T_ or π) as measures of genetic diversity. We calculate these estimators as describe in Tajima 1989 [75].

### Characterization of the flexible genome

From assemblies, we annotated the genomes using the RAST pipeline [77] with default parameters. Predicted proteins were used as input for the OrthoFinder software [78] to predict orthologous gene families. These orthologous gene families were classified into three different categories: multiple gene families (>1 copy per genome, on average), single copy genes (exactly one copy per genome) and flexible genome (<1 copy per genome).

### Presence or absence of genes

As absence of a given gene in a genome could be an artifact of the assembly process, we confirmed the absence of each gene family using the raw reads (Fig S2). A representative catalogue of the flexible genome protein sequences was built using the cd-hit program with a 90% similarity threshold [79]. We used sequences of the catalogue as queries for a blastn search on raw reads. We considered a gene family to be present in a given genome if the average coverage of the gene was greater or equal to 1X. This coverage threshold allowed us to detect every single gene in the gene catalogue (S3 Fig) while observing no variability among the control strains. To calculate coverage, we summed the length of all reads matching a given query over a minimal length of 100 nucleotides and a minimal identity of 97%, and divided by the gene length.

### Inference of flexible gene origins

In order to estimate the origin of the flexible genome gene pool, we performed an extended phylogenetic analysis of all 155 flexible gene families. The flexible gene catalogue served as query for a blastp search against the NCBI database. For each gene, we selected the first 200 hits matching with an E-value below 1E-05. The hit sequences were aligned using muscle with default parameters [80] and phylogenetic analysis performed with the FastTree algorithm using default parameters [81]. Finally, we screened nexus formatted trees from FastTree to identify the closest relative sequences of each gene in our dataset. This allowed us to classify the flexible genes into three mutually exclusive categories: first, genes whose closest relative originated from the *Vibrio cholerae* gene pool; second, genes whose closest relative is from *Vibrio* but not *cholerae* (i.e non-cholera *Vibrio* strains); and finally, the third category includes genes whose closest relative was outside the genus *Vibrio*. Trees were also displayed automatically using the FigTree java program for a manual inspection (http://tree.bio.ed.ac.uk/software/figtree/). To guard against false-positive inference of horizontal gene transfers from non-*V. cholerae*, a negative control was performed. We repeated the blastp/FastTree procedure using 155 genes extracted randomly from core genes in our study. As expected, these were all assigned *Vibrio cholerae* taxonomic affiliations.

### Phylogenetic evolutionary inference and root-to-tip regression

For all phylogenetic analyses, we did not consider the Integrative Conjugative Element (ICE), as the evolution of this region, a mutation hotspot, mostly reflects recombination events (S1 Fig). The ICE was defined as the region of MJ1236 chromosome 1 located between positions 87776 and 193789 (after reverse-complementation of MJ1236), according to a previous study [82]. A final alignment of 201 concatenated hqSNPs was generated from the core genome of the 35 genotypes (TC01-TC21 plus one to four unique genotypes per patient) and used for phylogenetic analysis. We used Seaview v.4.5.4 [83] to generate a maximum-likelihood phylogeny, employing a general time reversible (GTR) substitution model with four rate classes and SPR branch-swapping. All sites being variable in the alignment, we did not consider the proportion of invariable sites. To measure the extent of clock-like structure in our data set, we performed a linear regression of root-to-tip distances against dates of isolation, using the TempEst software [84]. Substitution rates and divergence times were then estimated using BEAST package v.1.8.3 [40], with XML-input files manually modified to specify the number of constant sites. For this analysis, we tested and compared both strict and uncorrelated lognormal molecular clock models and three coalescent models (exponential growth coalescent, constant-size coalescent and Bayesian Skyline demographic models), resulting in 6 possible model combinations. For all of them we used a GTR + G nucleotide substitution model and the sampling times for calibration. All of these combinations were run using 10,000,000 MCMC chains, with 10% burn-in and sampling every 5,000 generations. We used Tracer v.1.6 to ensure proper mixing, with all parameters having an effective sample size > 200. To select the best molecular clock and coalescent models, we estimate the marginal likelihoods for each combination via path-sampling, and we compared them with Bayes factors [85]. Divergence time, substitution rates and resulting tree were reported from the models with the highest marginal likelihood.

### Tests for natural selection over a three-year period

To distinguish between positive selection, purifying selection, or neutral evolution of protein-coding sequences, we considered variation in the proportion of nonsynonymous hqSNPs (*p*_NS_) in the *V. cholerae* core genome. Specifically, we evaluated how *p*_NS_ varied over time (a three-year period from 2011 to 2013) and among branches of the phylogenetic tree. We considered *N*=136 hqSNPs (excluding the ICE region, a mutation hotspot; S1 Fig) that varied among 21 isolates sampled over three years in Bangladesh (patients TC01-TC21) and the 122 isolates sampled from five patients from Bangladesh (patients B1-B5) and three from Haiti (Patient H1-H3). For these analyses, we excluded iSNVs by considering only the most frequent haplotype found within each patient, assumed to be ancestral.

We first tested whether the overall observed *p*_NS_ is to be expected under a simple neutral model of evolution. We performed 1,000 simulations of *N* mutations randomly distributed across the core genome of the MJ1236 reference. For each simulation, we re-estimated the relative proportions of intergenic (*p*_I_), synonymous (*p*_S_) and nonsynonymous (*p*_NS_) mutations, using annotations available for MJ1236 (GenBank: CP001485-6). We controlled for potential effects of genome-wide nucleotide composition by comparing simulations with or without imposed GC content of mutated positions (i.e. matching the GC content observed among the *N* real hqSNPs). We also controlled for any bias in the transition:transversion ratio by comparing simulations with or without imposed transition rate (i.e. matching the ratio observed among the *N* real hqSNPs). We considered that the core genome evolved under positive selection when the observed *p*_NS_ was higher than in at least 97.5% of simulations. We considered that the core genome evolved under purifying selection when the observed *p*_NS_ was lower than in at least 97.5% of simulations. We failed to reject neutral evolution when the observed *p*_NS_ fell within the 95% range of simulations.

Second, we tested whether fixed core genome hqSNPs were distributed evenly across branches of the phylogeny. Specifically, we asked whether substitution rates differ between Bangladesh and Haiti, or between long internal branches and the shorter, more recent tips of the tree where selection may have had insufficient time to act. To do so, we defined 3 well-separated monophyletic clades based on the evolutionary tree of the *V. cholerae* core genome. The tree was built in MEGA5 using a maximum composite likelihood model [86] (S6B Fig). We distinguished hqSNPs that were fixed among clades (corresponding to long branches) from those that are variable within clades (the tips of the tree). We hypothesized that if differences in *p*_NS_ are observed between vs. within clades, this could suggest that selection or substitution rates vary among clades and over time. To test this, we performed 10,000 random permutations of hqSNPs among branches of the evolutionary tree, and for each permutation, we re-estimated *p*_I_, *p*_S_ and *p*_NS_ within clade. For each monophyletic clade, we considered that the substitution rate was higher than expected by chance when the observed *p*_NS_ was higher than in at least 97.5% of permutations. We considered that the substitution rate was lower than expected by chance when the observed *p*_NS_ was lower than in at least 97.5% of permutations. We failed to reject neutral evolution when the observed *p*_NS_ fell within the 95% range of permutations.

### Tests for natural selection within and between patients

To investigate the role of natural selection within versus between patients from Bangladesh and Haiti, we performed the McDonald-Kreitman test [42] to test the neutral hypothesis that nonsynonymous (NS) to synonymous (S) substitution ratios remained constant over evolutionary time (within vs. between hosts). Specifically, we computed the Fixation Index (equivalent to an odds ratio statistic) as the NS:S ratio between patients (fixed SNPs) divided by the NS:S ratio within patients (iSNVs). Significant deviations of the Fixation Index from neutral expectation were evaluated using Fisher’s exact test.

We then tested whether iSNVs are equally distributed among patients, and if any patient contained an excess (possibly due to positive or balancing selection) or a deficit (possibly due to efficient purifying selection) of NS iSNVs. To do so, we performed permutations of iSNVs among the eight patients and estimated expected iSNV frequencies (*F*) and *P*_NS_ per patient (B1-B5; H1-H3) and region (Bangladesh and Haiti). We first assigned each of the 122 strains collected from the eight patients to one of the four following haplotypes: H_0_ as strains having the most frequent haplotype found within each patient, and assumed to be ancestral for that patient; H_INT_ as strains having one intergenic iSNV; H_S_ as strains having one synonymous iSNV and H_NS_ as strains having one synonymous iSNV. (No strains were observed with more than one iSNV, so these haplotypes are sufficient to model the observed intra-patient diversity). We then performed 10,000 random permutations of the four haplotypes among the 122 strains. When an iSNV was shared between two or more strains within a patient, but not observed in other patients, we ensured that these strains were always assigned to the same patient during permutations. For each simulation and for each patient or region, we reported the total iSNV relative frequency (*F* = number of strains containing iSNVs / the total number of strains sequenced for that patient or region) and *p*_NS_, defined as above. For each region and each patient, we considered that *F* and *p*_NS_ were higher or lower than expected by chance when the observed values were respectively higher or lower than in at least 97.5% of permutations. We concluded that *F* and *p*_NS_ were consistent with our neutral model when they fell within the 95% range of permutations.

### Sensor histidine kinase protein conservation analysis

Of the three NS iSNVs in patient H1, two occur in the same gene, a predicted sensory histidine kinase (NCBI accession number ACQ61177, from the reference genome MJ-1236). We sought to determine whether these two NS iSNVs occurred in conserved or variable peptides. To do so, we retrieved the 500 best matches (top BLAST hits) for the ACQ61177 protein sequence in NCBI GenBank using BLASTp. From these 500 homologous sequences, we removed identical (duplicate) sequences, and those that were truncated at the N or the C terminus, resulting in 465 unique homologous sequences. We then determined whether 4-amino-acid (4aa) peptides surrounding the mutated residues were conserved among these sequences. We defined a simple conservation score as the proportion of homologs having the reference peptide (from *V. cholerae* MJ1236). This score could be influenced by a biased sample of sequences in GenBank, and thus represents a rough estimate of conservation. In order to minimize the effect of peptide convergence, we did not consider 4aa motifs that were found at least twice in at least one sequence, which was not the case for any of the peptide motifs affected by the two observed iSNVs.

### Liquid culture and biofilm growth assays of isolates from patients H1 and H2

We performed *in vitro* experiments to identify phenotypic consequences of iSNVs in patient H1, or flexible gene content variation in patient H2. Isolates were grown in 4 ml of LB broth with agitation at 30°C, and optical densities were measured at 600 nm using a spectrophotometer every hour for 12 hours, revealing no significant differences in growth between strains. To test for biofilm production, we grew the same strains in 200 μl of LB in a 96-well plate, without agitation at 30°C for 48 hours. Controls included empty wells, wells with LB only, and wells with a *V. cholerae* strain with an in-frame deletion in the *vpsA* (Vibrio polysaccharide A) gene, which results in reduction of biofilm production[47]. <u>A</u>0.1% solution of Crystal Violet was used to stain for biofilm adherent to the well. Biofilms were dissolved in ethanol at the end of the assay, and the optical density was measured at 595 nm using spectrophotometry. Experiments were performed in replicates of four to twelve.

### Polymyxin B MIC assay on patient H1 isolates

Strains H1C1, H1C5 and H1C6 were grown overnight at 30°C on LB agar and then cultures were diluted 1:100 in fresh LB medium. Cells were grown to midexponential growth and diluted 1:10, and an aliquot was plated on LB agar. Polymyxin B Etest gradient strips (AB Biodisk) were applied onto inoculated plates and incubated at 37°C, and the MICs were evaluated after 16 h.

## Acknowledgments

We thank Simone Perinet and Meri Debela for their technical assistance, Paula Watnick who contributed materials to the study, and Salvador Almagro-Moreno for constructive comments on the manuscript. We also thank the Zanmi Lasante staff at the enteric microbiology laboratory in St. Marc, Haiti. Finally, we are grateful to the people of Dhaka where our study was undertaken; to the field, laboratory and data management staff who provided tremendous effort to make the study successful; and to the people who provided valuable support in our study. ICDDR,B gratefully acknowledges the Government of the People’s Republic of Bangladesh; Global Affairs Canada (GAC); Swedish International Development Cooperation Agency (Sida) and the Department for International Development, (UKAid). We declare that we have no competing financial interest.

### Funding information

This study was supported by CIHR (Canadian Institutes of Health Research) and the Canada Research Chairs program, the ICDDR,B: Centre for Health and Population Research, grants AI099243 (J.B.H and L.C.I), AI103055 (J.B.H and F.Q), AI106878 (E.T.R and F.Q.), AI058935 (E.T.R, S.B.C and F.Q.), T32A1070611976 and K08AI123494 (A.A.W.) from the National Institutes of Health, and the Robert Wood Johnson Foundation Harold Amos Medical Faculty Development Program (R.C.C.).

